# The discrete logic of the Brain - Explicit modelling of Brain State durations in EEG and MEG

**DOI:** 10.1101/635300

**Authors:** Nelson J. Trujillo-Barreto, David Araya, Wael El-Deredy

## Abstract

We consider the detection and characterisation of *brain state* transitions, based on ongoing Magneto and Electroencephalography (M/EEG). Here a brain state represents a specific brain dynamical regime or mode of operation, which produces a characteristic quasi-stable pattern of activity at topography, sources or network levels. These states and their transitions over time can reflect fundamental computational properties of the brain, shaping human behaviour and brain function. The Hidden Markov Model (HMM) has emerged as a useful model-based approach for uncovering the hidden dynamics of brain state transitions based on observed data. However, the Geometric distribution of state duration (dwell time) implicit in HMM places highest probability on very short durations, which makes it inappropriate for the accurate modelling of brain states in M/EEG. We propose using Hidden Semi Markov Models (HSMM), a generalisation of HMM that models the brain state duration distribution explicitly. We present a Bayesian formulation of HSMM and use the Variational Bayes framework to efficiently estimate the HSMM parameters, the number of brain states and select among alternative brain state duration distributions. We assess HSMM performance against HMM on simulated data and demonstrate that the accurate modelling of state duration is paramount for accurately and robustly modelling non-Markovian EEG brain state features. Finally, we used actual resting-state EEG data to demonstrate the approach in practice and conclude that it provides a flexible parameterised framework that permits closer interrogation of possible generative mechanisms.

## 1 Introduction

There is great current interest in the identification and characterisation of short lived and recurrent *Brain States (BSs)* underlying the non-stationary resting-state (and task evoked) Magneto/Electroencephalography (M/EEG) data (Khanna et al., 2015; O’Neill et al., 2018). Here a BS represents a brain dynamical regime or mode of operation producing a quasi-stable pattern of activity at topography, sources or network levels. These states and their transitions over time can reflect fundamental computational properties of the brain, shaping human behaviour and brain function (Ritter et al., 2015). The above inference task amounts to solving the so called BS allocation problem (Olier et al., 2013; Woolrich et al., 2013), which is challenging. The single most important challenge is that the BS and its temporal dynamics are not observable, but live in a latent (hidden) space the structure which we also have to uncover. The unknown structure includes the number of brain states and the specific characteristics of the M/EEG features associated with a particular BS. An accurate *Brain Sate* allocation therefore requires making realistic assumptions about the dynamic generative mechanisms producing the observed fingerprints on data. These mechanisms are still not well understood, and therefore the required approach must also offer a principled way to evaluate the suitability of the assumptions made.

Given their high temporal resolution, M/EEG is ideal to interrogate the short temporal scale at which BS changes occur (millisecond range) (O’Neill et al., 2018). Slow fluctuations in the power of EEG frequency bands have been found to correlate with temporal changes of resting-state networks in functional Magnetic Resonance Imaging, occurring in the hundreds milliseconds to seconds time scale (Preti et al., 2017). However, M/EEG has also provided strong evidence that the brain can switch between BSs at a significantly faster rate with average BS durations in the order of tens to a few hundred milliseconds (Baker et al., 2014; Cabral et al., 2014; Khanna et al., 2015; Michel and Koenig, 2018; Olier et al., 2013; Vidaurre et al., 2016; Woolrich et al., 2013). More importantly, the observed covariation with the slower RSNs in fMRI has been proposed to be mediated by underlying dynamic mechanisms endowing M/EEG extracted BS sequences with long-term dependencies and scale-invariance extending across several temporal scales (Cocchi et al., 2017; Gschwind et al., 2015; Roberts et al., 2015; Van De Ville et al., 2010). Moreover, important information about these dynamic mechanisms is carried in the BS switching times, with BS duration distributions being predicted to be heavy-tailed (Roberts et al., 2015).

Existing methods to approach the BS allocation problem has been dominated by data-based approaches, including temporal sliding windows (Hansen et al., 2015; O’Neill et al., 2017), adaptive segmentations using clustering (Hassan et al., 2015; Khanna et al., 2015; Mheich et al., 2015), among others (O’Neill et al., 2018; Preti et al., 2017). Although the shortcomings of these approaches are well stablished (Hindriks et al., 2016; O’Neill et al., 2018), from a modelling perspective, their main limitation is that the temporal dynamics of BSs is not considered during estimation, but it is assessed a posteriori.

Model-based approaches consider the BS allocation problem based on a generative (parametric) model summarising our beliefs of how the hidden BS sequence gives rise to the observed M/EEG data features (Olier et al., 2013). The inversion of this model based on data allows for the inference of model parameters, consistent with the model assumptions. The recurrent behaviour of BSs suggests a Hidden Markov Model (HMM) approach which has proved to be a useful modelling tool for M/EEG BS allocation (Baker et al., 2014; Vidaurre et al., 2018, 2016; Woolrich et al., 2013). In brief, the HMM uses a Markov Chain to model the transitions between a finite set of hidden discrete BSs, so that each time a BS is active it emits an observation. However, the Markovian assumption implicit in HMM implies that the distribution of BS duration is strictly Geometric, thus favouring BSs with short durations. Therefore, using HMM to model M/EEG data with long-term dependencies and persistent BSs (i.e. non-Markovian behaviour) will allocate high probability on BS sequences with unrealistically fast switching. Therefore while standard HMM seems like a natural fit to the BS allocation problem in M/EEG, it inadequately models the temporal properties of observed M/EEG time-series (Cocchi et al., 2017; Roberts et al., 2015; Van De Ville et al., 2010; von Wegner et al., 2017).

This key limitation to the HMM motivates our investigation to extend it into a more flexible model called Hidden Semi Markov Model (HSMM) (Yu, 2010). HSMM is a generalisation of HMM where the Markovian assumption is relaxed to allow for explicit modelling of the BS duration distribution. The duration model can naturally encapsulate prior beliefs that BSs are long, while at the same time allowing BS recurrence. This flexibility comes at the cost of having to consider additional parameters for the state durations. We will show however that, in the context of BS allocation for M/EEG, the added complexity is more than compensated for by the explanatory power of the HSMM model. We demonstrate that HSMM can account for most of the limitations of HMM in the BS allocation setting. We use a Variational Bayes formulation of HSMM (Hudson, 2009), which allows for coherent and efficient estimation of the hidden BSs sequence, its parameters and its dimensionality (number of BSs) even at the level of a single subject. More importantly, the proposed Bayesian framework offers a principled way of choosing among alternative BS duration distributions.

The paper is organised as follows. In sections 2.1–2.4 the technical aspects of HSMM are described highlighting its main differences with respect to the standard HMM. Sections 2.5–2.7 present the computational simulation framework and the empirical data used to evaluate HSMM performance in relation to HMM. Sections 3 and 4 describe respectively the results obtained and a discussion of the main findings in relation to the state-of-the-art of the literature. A MATLAB (The Mathworks, Inc) implementation of our Brain State Dynamics (BSD) toolbox is also available (https://github.com/daraya78/BSD).

## 2 Methods

In this section, we describe the proposed HSMM for BS allocation based on ongoing M/EEG data and highlight the key differences with the standard HMM approach. The model is motivated by Hybrid Dynamical System (HDS) theory for Multivariate Time Series Analysis applied to M/EEG. We summarise the framework and refer interested readers to supplementary material and previous descriptions of these models for additional mathematical and algorithmic details.

### 2.1 Hidden Semi Markov Model for dynamic Brain State allocation

HSMM and HMM are successful models of HDS. Dynamic Bayesian Network (DBN) representations of these models are shown in Figure 1. A HDS produces continuous multivariate time-series data, but its dynamics can rapidly transition among a discrete set of states or modes of operation. HSMM and HMM capture such dual behaviour using two stochastic processes. The underlying process describes the hidden sequence of discrete states with its own dynamical evolution. This influences the continuous process that generates the data or emissions (Rabiner, 1989; Yu, 2015).

**Figure 1.**
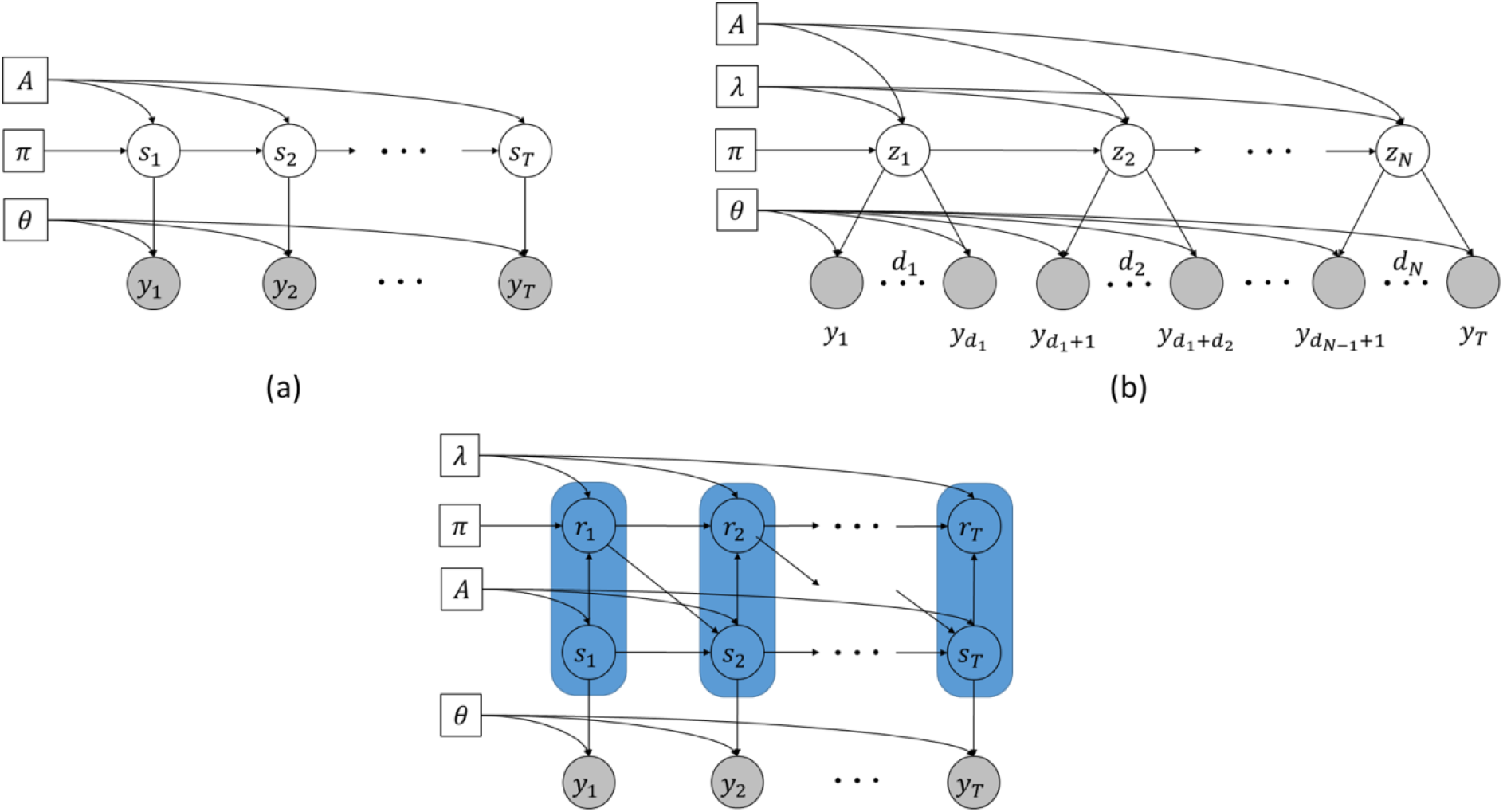
Dynamic Bayesian Network (DBN) representations of the hybrid system models used in this paper. Shaded circles represent observations, clear circles represent hidden state variables and squares represent parameters. (a) Standard Hidden Markov Model (HMM): One observation is admitted from the active state at each time point. (b) HSMM interpreted as a hidden standard Markov chain on a set of “super-state” nodes 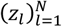. The super-states emit random length segments of observations, of which the first *T* are observed. (c) HSMM interpreted as a hidden augmented joint-state process 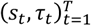 of states and residual times. The augmented joint-state is highlighted in blue. Note that only one observation per realisation of the joint-state is emitted at each time point. When first entered the state, the residual time is value sampled from a duration distribution; it then deterministically counts down to *r_t_* = 1 at which point the state is free to change and ***r_t_*** is set to the duration of the new state.

HSMM and HMM however differ in the way state durations are considered. In HMM, the hidden process is a Markov chain where *only one* observation per state is emitted at each time point (Figure 1A). Therefore, in HMM a state’s duration is implicitly captured as self-transitions (transitions to the same state), which implies a Geometric distribution over state’s duration (Rabiner, 1989). In HSMM the hidden process is a semi-Markov chain where each state is a “super-state” that can emit a segment of observations with variable length defining the duration of the state (Figure 1B). There are several modelling approaches to semi-Markovianity (Murphy, 2002; Yu, 2010), but here we focus in the setting where each state’s duration is given an explicit probability distribution function.

The HSMM has four components: the initial distribution, the transition distribution, the duration distribution and the emission distribution of the BSs. Let’s denote the observed M/EEG feature sequence of length *T* at *n* emission channels as the *n*-variate time-series *y*_1:*T*_ (*y_t_* ∈ ℝ^*n*^). Let’s also consider the equivalent representation of the HSMM shown in Figure 1C in terms of the time-resolved hidden BS sequence *s*_1:*T*_ and the corresponding sequence of BS residual times *r*_1:*T*_. The BS *s_t_* at time *t* can take values in the set 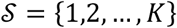 of *K* possible brain modes (*s_t_* ∈ *S*); while *r_t_* takes values in the set 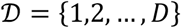 of *D* possible BS durations 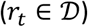. This representation renders the HSMM equivalent to a homogeneous HMM, by considering the augmented joint process (*s_t_,r_t_*) as a single Markov Chain (Hudson, 2009) in the joint space 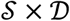, with a general transition kernel

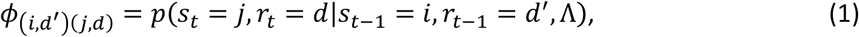

Where Λ are parameters of the transition distribution of the joint process. The advantage of the augmented joint-state over the super-state representation is that the Variational Bayes (VB) inference algorithms developed for standard stationary HMM can be employed directly.

The HSMM used here assumes that (i) transitions between states only occur at the end of the segment at which point (ii) self-transition is not permitted; (iii) the duration distribution generates a segment length at every state switch; and (iv) the segment length only depends on the generating state. Under these assumptions, the general transition kernel (1) takes the form

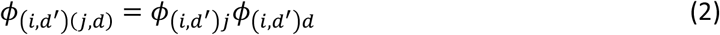

where 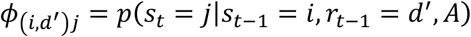 and 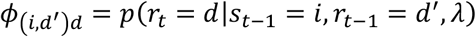 so that

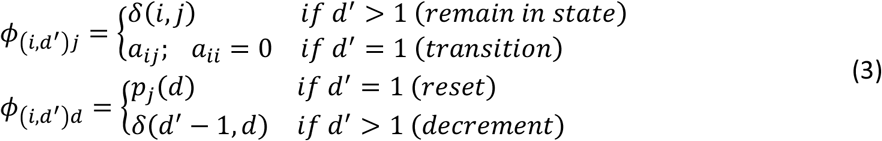

where the Kronecker’s delta *δ*(*i,j*) = 1 if *i* = *j* and zero otherwise. We have used the notation *p*(*s_t_* = *j*|*s*_*t*−1_ = *i*, *r*_*t*−1_ = 1, *A*) = *a_ij_* for all *t* (homogeneous Markov process), where *A* is the transition matrix of size *K* × *K* with elements *a_ij_*; and *p*(*r_t_* = *d*|*s_t_* = *j*, *r*_*t*−1_ = 1, *λ*^(*j*)^) = *P_j_*(*d*), where *λ*^(*j*)^ are the parameters of the duration distribution of state *j*.

Equation (3) means that if the brain transitions from mode *i* to mode *j* at time *t* with probability *a_ij_*, then it will remain in mode *j* for a random duration *d* sampled from the distribution *P_j_*(*d*), while the residual time deterministically counts down from *d* to 1. At the end of the segment, the brain is allowed to transition to a new state with a new duration and the residual time is reset to the new duration *d*.

We modelled durations using a truncated Normal distribution in simulated data

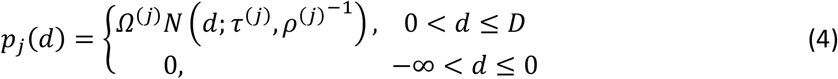

and compare this to using a truncated Log-normal distribution in the empirical data

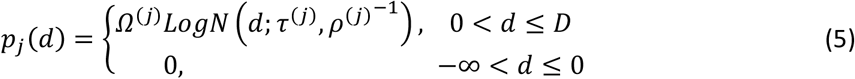

Where *λ*^(*j*)^ = [*τ*^(*j*)^,*ρ*^(j)^] are the parameters of the duration distribution of the *j*-th BS and *Ω*^(*j*)^ is a normalizing factor so that 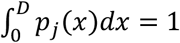. Other options of duration distributions are included in our publicly available MATLAB Toolbox.

Since our focus is on the adequate modelling of BS durations, without loss of generality we assume that the BSs are characterized by simple features where the emissions from the *k*-th BS are independently sampled using a multivariate Normal distribution of dimension *n* with state-specific mean *μ*^(*k*)^ and precision *Σ*^(*k*)^ (Baker et al., 2014; Hunyadi et al., 2019; Rukat et al., 2016; Woolrich et al., 2013). The conditional probability of emitting an observation *y_t_* given that the BS was in mode *k* at time *t* is then

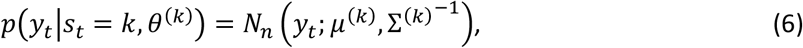

where 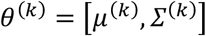

The initial distribution *π* = {*π_kd_*} is defined so that

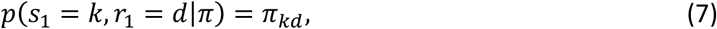

with *k* = 1,…, *K* and *d* = 1, …,*D*, and represents the probability of the *k*-th BS being active with residual time *d* at the beginning of the sequence. We adopt a usual simplification of this initial condition by assuming that the observations begin at a segment boundary, so that *r*_1_ = 1 and *π_kd_* = 0 for *d* ≠ 1. To completely define a HSMM, boundary conditions at the end of the observation sequence most also be defined. A common assumption is that the observation sequence also ends exactly on a segment’s boundary. We used a more realistic assumption that the observations are censored at the end, so that the final segment may possibly be cut off in the observations (see (Guédon, 2007) for details and alternative conventions)

### 2.2 Bayesian formulation and inference

We followed the Variational Bayes (VB) approach to HSMM in (Hudson, 2009), and adapted it to the model specification used here. The specification of the HSMM defines a complete likelihood function of the form

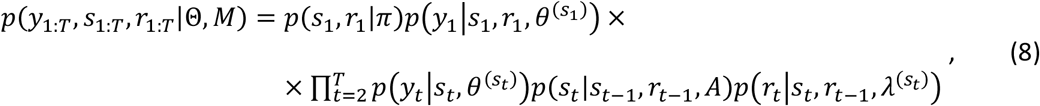

where *θ* = *θ*^(1:*K*)^ and *λ* = *λ*^(1:*K*)^ denote the collection of parameters of the emission and duration distributions of all the states, respectively; *M* is the model class defined by the choice of the model type (HSMM or HMM) and/or by the choice of the number of BSs given a model type; and Θ = {*π,A,θ,λ*} are the parameters which specify models within the class. Note that the HMM is completely defined by the reduced sets of parameters Θ = {*π,A,θ*}, because the duration distribution is not explicitly defined for that model. We have omitted *M* in the right-hand-side of (8) for notation simplicity.

The Bayesian approach requires prior distributions to be defined on all the parameters of the model. After combination with the likelihood via the Bayes Rule, a posterior distribution of the parameters and the hidden variables given the data is obtained, based on which statistical inference can proceed. We assume that all parameters are independent a priori so that the posterior is:

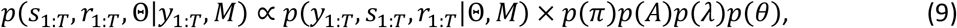

We further assume conjugate prior distributions for all the parameters (see Supplements).

Exact Bayesian inference based on (9) is not possible due to the dependence between the parameters and the hidden states. However, the augmented state representation of HSMM allows using the efficient VB algorithm developed for HMM (Beal, 2003; Hudson, 2009). VB algorithm for HSMM allows making inference on parameters, hidden states and models by approximating the joint posterior of hidden states and parameters, with a simpler variational density. The usual Mean Field Approximation (Ghahramani et al., 2000) requires the approximate posterior to factorise over subsets of parameters and hidden variables:

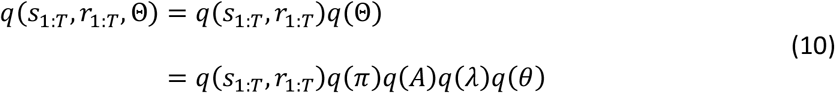

Note that the structure of the HSMM implies an exact posterior that is already factorised over groups of parameters (equations (8) and (9)), therefore only the factorisation between the parameters and the hidden states is an actual approximation in (10). In the present paper additional factorisations were required over the parameters defining the duration, and the emission distributions (Supplements).

The VB algorithm then aims to minimize the Kullback-Leibler (KL) divergence (Kullback and Leibler, 1951) between the true and the approximate posterior. This is equivalent to maximizing a lower bound on the logarithm of the model’s evidence called the Negative Free Energy (NFE). By approximating the model’s evidence, the NFE becomes a measure of model quality that trades model accuracy for model complexity and can be used for both model class selection and for monitoring the convergence of the VB algorithm. The Supplements contain mathematical details of the VB framework specific to the models’ setup presented here.

### 2.3 Procedure for estimating the number of BS

The number of BS is a structural parameter, therefore inferring its optimal value given the data, amounts to a model class selection task. In the VB framework proposed here, we used a selection strategy based on the NFE, similar to (Olier et al., 2013). This strategy entails running the VB algorithm multiple times, each with a different initial number of BSs. In each VB run, an implicit Automatic Relevance Determination procedure was also implemented, whereby BSs which did not receive support from the data were automatically removed from the analysis. The nuisance BSs were defined as those getting negligible probability of being active at all time points. The optimal number of BSs was the one with maximal NFE across all runs.

### 2.4 Simulation framework

We used computational simulations as a test ground for evaluating the performance of HSMM vs HMM for BS allocation. The simulations’ pipeline is shown in Figure 2. All simulations were based on 3 BSs, each characterized by a specific EEG topography map and a specific data covariance matrix. The BS maps were obtained using a generative process, whereby a simulated brain source activity was projected to the sensor space using the linear EEG Forward Model (Trujillo-Barreto et al., 2004). A realistic lead field matrix relating the source and sensor spaces was computed based on the digital brain phantom developed at the Montreal Neurological Institute (MNI) (Collins et al., 1998). The source space consisted of a mesh of *G* = 20000 current dipoles located at the vertices of the tessellated grey/white matter interface. The sensor space consisted of the standard 128 electrodes BioSemi system, which was digitally placed on the scalp of the phantom. In each simulation experiment, the source activity image associated with each BS was sampled from a G-variate Normal distribution with zero mean and covariance matrix *∑_j_* = (*L^T^L*)^−1^, where *L* is the discrete surface Laplacian operator (Huiskamp, 1991) defined on the nodes of the vertices of the source space. This choice of *∑_j_* is consistent with the assumptions of the popular LORETA source reconstruction method (Pascual-Marqui et al., 1994) and ensured producing spatially smooth source activity. After projection to the sensor space, the EEG map associated with each BS were normalised to have unit norm.

**Figure 2.**
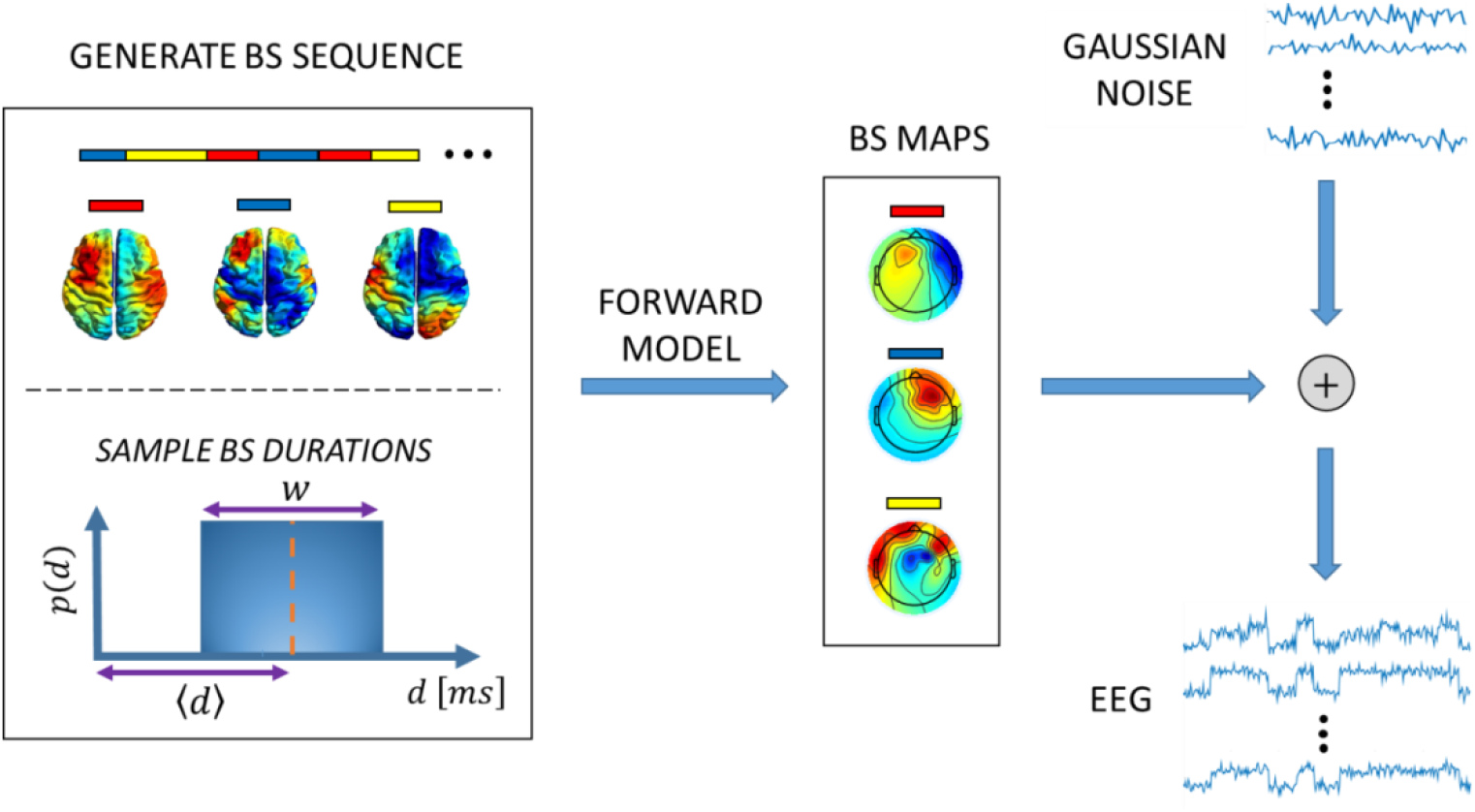
Simulation pipeline. EEG activity is simulated considering the brain as a Hybrid Dynamical System. The hidden discrete state process represents source activations that remain stable over a time segment before rapidly switching to a different source activation. The length (duration) of the time segments are randomly sampled from uniform distributions of durations with state-specific mean 〈***d***〉 and width ***w***. The source activations are instantaneously projected to the sensors’ space using the linear EEG Forward model and subsequently corrupted with noise to produce the observed EEG activity.

The temporal dynamics of the EEG signals where generated considering the brain as a HDS where, at each BS, the brain emitted a segment of observations of length sampled from a uniform duration distribution centred on a BS-specific mean duration 〈*d*〈 and with a BS-specific width *w*. This simple duration model allowed for generating BS sequences with different memory (temporal dependence) lengths. During the life time of a BS, observations were sampled from a multivariate normal distribution with constant mean and diagonal covariance matrix. The total length of the observed data in each realisation of a simulation experiment was *T* = 4000 in all cases. The mean of the BS emissions was the scalp-projected source activity associated with the BS, and the covariance matrix modelled the (possibly BS-specific) zero mean Gaussian iid observation noise. The variance of the noise was chosen to achieve a fix Signal-to-Noise Ratio (SNR) across data sequences of the same simulation experiment (Appendix A).

BSs’ transitions were also designed to emulate non-Markovian properties of empirically observed EEG microstate sequences (von Wegner et al., 2017). That is, at the end of a BS segment, a new BS was randomly generated based on the asymmetric transition matrix:

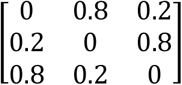

This choice of matrix favours cyclic BS transitions (*BS*1 → *BS*2 → *BS*3). Without loss of generality, in all simulations the system was assumed to be in the first BS at the beginning of the sequence.

Since our focus is on the modelling of BS durations, BS maps were generated ensuring a fair degree of dissimilarity between the maps of different BSs. This was done so that differences in performance found between the evaluated methods were due to the differences in the way BS duration is modelled, rather than due to inability to discriminate between similar BS maps. Results of more extensive simulations assessing the effect of similarity between the BS maps were found to reinforce the conclusions of the paper, and can be provided on request.

### 2.5 Scores used to evaluate model performance

The mathematical expressions of the evaluation scores used are summarised in Appendix B. First, following (von Wegner et al., 2017), the Auto Information Function (AIF) was used to demonstrate that the HSMM was capable of producing dynamical features observed in the well-studied EEG microstate sequences. By analogy with the Autocorrelation Function for real valued signals, the AIF denotes the time-lagged mutual information between the BS signal *s_t_* and delayed versions of itself (*s_t+l_*). It measures the amount of information about *s_t+l_* contained in *s_t_*. Then, the performance of the proposed HSMM was compared to the standard HMM in terms of their suitability as a generative model for BS allocation The Sequence Accuracy Score (SAS) measured the similarity between the decoded and the simulated BS sequences. SAS was defined as a function complementary to the Hamming Distance (Navarro, 2001) between the two sequences, expressed in percentage. Given two sequences of equal length, the Hamming Distance is an edit distance defined as the minimum number of substitutions required to change one sequence into the other. SAS is then defined so that a value of 100% means the two sequences are identical (no substitutions needed), while 0% indicates no matching between any of the elements of the two sequences (all elements need to be substituted). Additionally, the Jensen-Shannon Divergence (*JSD*) (Endres and Schindelin, 2003) was used to measure the dissimilarity between the histograms of BS durations of the simulated sequence and the estimated one.

We also used model quality measures to evaluate the relative performance of two models (or two model classes) based on information theoretic principles, which do not require comparing to a gold standard. The Log-Bayes Factor *logBF*(*M*_1_, *M*_0_) (Kass and Raftery, 1995) was used to measure the evidence provided by the data in favour of the model class *M*_1_ (e.g. HSMM) against *M*_0_ (e.g. HMM). Additionally, we used a measure of Discriminative Power Score *DSP*(*H*_1_,*H*_0_) (Juang and Rabiner, 1985) to evaluate the ease of discriminating between two models given a data sequence. That is, if H0 and H1 are the hypotheses that the data sequence was generated by a certain HSMM with parameters *Θ*_0_, or another HSMM with parameters 0_1_, respectively, then *DSP*(*H*_1_, *H*_0_) represents the average information per data sample for discrimination in favour of H1 against H0.

### 2.6 Empirical data acquisition and analysis

Ten minutes of spontaneous task-free EEG were recorded from one healthy subject. The subject was asked to seat comfortably with eyes open during the experimental session. The EEG data were recorded from 64 scalp electrodes using the ActiveTwo system (BioSemi, Amsterdam, Netherlands) and Actiview^®^ acquisition software (Biosemi Netherlands) at a sampling rate of 2048Hz.

#### Data pre-processing

For consistent comparisons, we followed a similar pre-processing pipeline as in (Baker et al., 2014), but slightly modified to the case of EEG data. The pipeline was implemented in MATLAB as follows. Channels and periods of data containing obvious artefacts were visually identified and discarded (5%). The signals were band-pass filtered between 0.1Hz and 120Hz using a high-pass and low-pass FIR filter in sequence; and were subsequently de-trended and down-sampled to 512Hz. Artefact detection and correction was carried out by decomposing the data into 30 temporally Independent Components using Independent Component Analysis. Stereotyped components related to ocular, cardiac, motion, and mains interference artefacts were classified and rejected based on their topographic, temporal and spectral features. The remaining components were then used to reconstruct the clean data. Bad channels were interpolated using spline interpolation and the data was subsequently re-referenced to average reference for further analysis.

#### Data preparation for HSMM analysis

Following previous studies, the HSMM was applied to the power envelopes of the EEG signals (Baker et al., 2014; Hunyadi et al., 2019; Rukat et al., 2016). The pre-processed data was first band pass (FIR) filtered between 4 and 30Hz. The amplitude envelope of the oscillatory activity at each channel was then derived by computing the magnitude of the analytic signal, obtained from the Hilbert transformed data. For computational efficiency, the envelope amplitudes were down-sampled to 40Hz using a polyphase anti-aliasing filter as implemented in the MATLAB signal processing toolbox. Principal Component Analysis was then used on the normalised envelopes (zero mean and unit standard deviation) for whitening and dimension reduction to keep the first 40 principal components (PC) (99.21% explained variance).

#### HSMM setup

Two HSMMs were inferred using either a Normal or a Log-Normal duration distribution. In all models, the BSs were assumed to emit observations using each a 40-variate Normal distribution over the observations (here principal components of the envelope data) specified by the corresponding mean vector and covariance matrix.

#### BS topographic maps

Due to the dimensionality reduction via Principal Component Analysis, the means and covariance matrices characterising each BS are not straightforwardly interpretable since they are estimated in the low dimensional space spanned by the principal components. To obtain interpretable scalp activity maps associated with each BS, we followed a procedure similar to (Baker et al., 2014). We computed the partial correlation of the BS time courses with the amplitude envelope data at all the electrodes. The obtained correlation maps represent the unique contribution of each BS to the oscillatory activity power, while components of the envelope signals that are common across BSs are minimised.

The partial correlation was computed using a general linear model analysis (Friston et al., 1996) with the BS signals as predictors and the full-rank envelop data (prior to dimension reduction and whitening) as response variables. Each BS signal was a binary vector indicating whether the BS is on or off at each point of the most probable BS path, which was estimated using the Viterbi algorithm for HSMM (Yu, 2015). Given the estimated Viterbi path *Ŝ*_1:*T*_’, each element of the [*T* × *K*] design matrix is

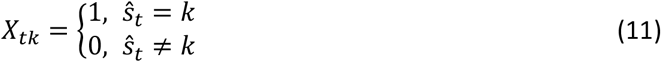

The columns of *X* and the envelope data were both z-scored. The *K* vectors of estimated regression coefficients defined scalp spatial maps representing estimates of the partial correlation coefficient between each BS and the envelope data. In order to facilitate interpretation, after inference the BSs were re-labelled so that BSs associated with similar maps across models, were assigned the same label.

#### 2.7 Initialisation of the VB algorithm

To account for possible dependence of the inference on the initial conditions (e.g. convergence to local minima), both the HSMM and the HMM VB algorithms were ran 10 times each using different starting points. Each starting point was determined by first running a k-means clustering algorithm in order to produce a BS sequence by assigning each data point to the closest k-mean cluster. The k-means algorithm was itself initialised 3 times using random data points as initial centroid positions and the solution producing the most compact clusters was chose as the optimal one. The sample estimates of the models’ parameters obtained based on the k-means BS sequence were then used as initial values for the VB iterations. The optimal VB initialisation was determined as the one with highest NFE after 10 iterations of the VB algorithm. In practice we found this procedure to consistently produce identical results as compared to running the VB fully until convergence several times and then choosing the solution with highest NFE, but it was significantly more efficient computationally.

## 3 Results

### 3.1 Data simulation test

We first demonstrated that the semi-Markov chain embedded in our HDS simulator can reproduce key dynamical properties of empirically observed EEG microstates sequences. These properties are consistent with non-Markovian behaviour (von Wegner and Laufs, 2018) including: (i) non-Geometric duration distributions; (ii) non-symmetric and cycling transitions; (iii) non-stationary transitions in the Markov sense (i.e. considering transitions at every time point); and (iv) relatively long but finite memory (temporal dependencies) demonstrated via distinct periodicities of the AIF of microstate sequences, with a tail that converges asymptotically to Markovian memoryless in the long run. Since properties (i)-(iii) are ensured by the definition of the HSMM, we focus on property (iv).

Figure 3 shows the AIF surfaces of sequences generated from semi-Markov and Markov chains. The sequences were generated using HSMM and HMM without emissions. The HSMM used 3 BSs with Uniform durations and the transition matrix defined in section 2.4. For each sequence, the mean durations (Figure 3a,c) or widths (Figure 3b,d) of the 3 BSs were varied at once. The HMM sequence matching each HSMM sequence was generated using the empirical transition matrix calculated based on the corresponding HSMM sequence. The AIF of HSMM showed distinct oscillations with amplitude decaying slowly with the time lag. This is consistent with the behaviour of EEG microstates sequences obtained using empirical data (von Wegner et al., 2017) and corresponds to a process with memory (time dependencies), but Markovian “memoryless” in the long run. Interestingly, the AIF peaks were located at lags which were multiples of the BS mean duration and changes in the position of the peaks followed the changes in the mean duration. Additionally, the amplitude of the AIF peaks was inversely related to the mean duration. Changes in the width however, only affected the amplitude of the AIF oscillations, so that increased widths were associated with reduced amplitudes. The AIF of the Markov chain showed a featureless fast and monotonic decay with the time lag in all cases. The decay rate was higher for smaller mean duration values, while it was insensitive to changes in the width of the duration distribution.

**Figure 3.**
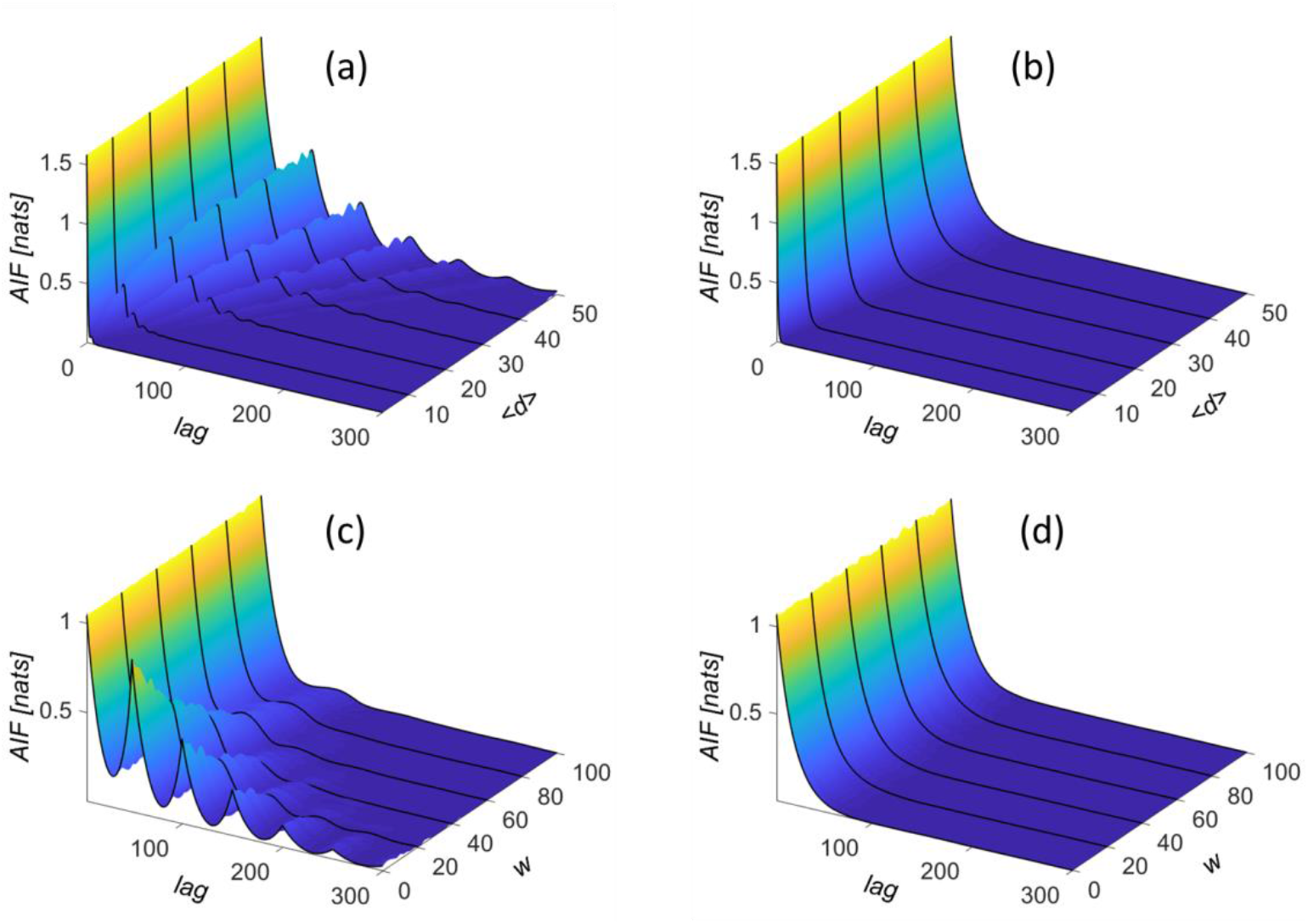
Reproducing microstates features in simulated BS sequences. Auto Information Function (AIF) of the BS sequences generated using HSMM (a & c) and HMM (b & d) without emissions, as a function of the mean and width of the duration distributions of the BSs. In (a) and (b), the mean duration of the three BSs was varied while the widths were fix at 6 a.u. In (c) and (d), the widths were varied while the duration means were fix at 50 a.u. The periodicities and the asymptotic decay in the case of the HSMM resembles that of the empirically observed EEG microstates, as illustrated by (von Wegner et al., 2017). The AIF is measured in natural units of information (nats).

A reproducibility study was then carried out to investigate whether good data fitting was a sufficient criterion for a HSMM or a HMM to reproduce the underlying BS dynamics of a non-Markovian HDS. HSMM and HMM were both trained on simulated EEG datasets (see section 2.4) with different memory lengths of the underlying BS sequence. The memory length was varied by manipulating the mean (centre) of the uniform duration distribution of a test BS. The duration distributions of the other two BSs had fixed mean (50 a.u.) and width (6 a.u.) across all simulations. After training, new BS sequences were generated from each of the trained models and the resultant histograms of durations corresponding to the test BS were compared to the actual sampling duration distribution that generated the training dataset.

Figure 4 shows one exemplary histogram. Although the two models had an excellent goodness-of-fit in the training phase (not shown), they performed quite differently in their ability to reproduce the hidden dynamics of the HDS. As expected, the geometric distribution implicit in HMM allocated nonzero probability mass across the whole range of possible duration values, while the explicit normal duration distribution in HSMM concentrates its probability mass in the correct range of duration values.

**Figure 4.**
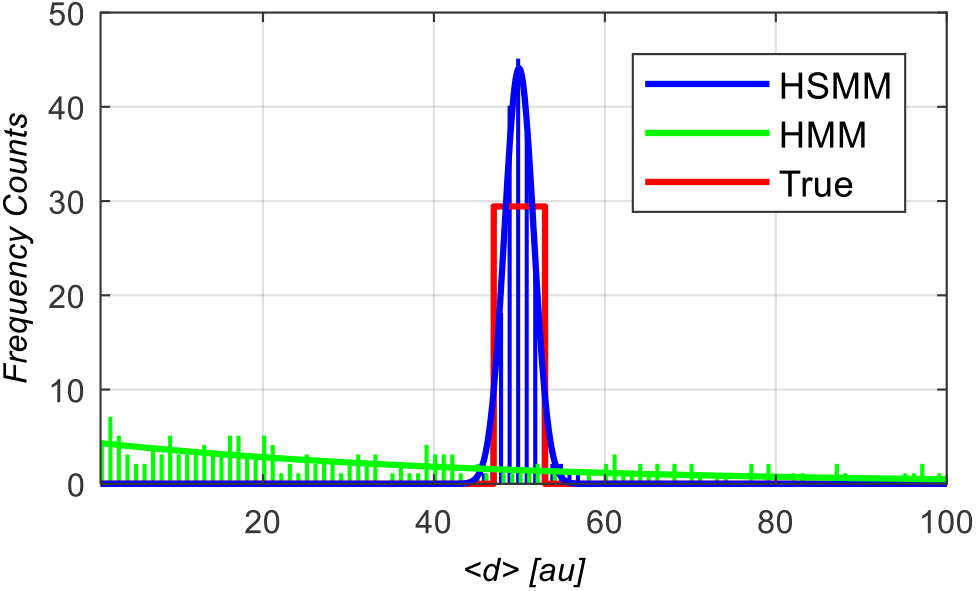
Histograms of durations of a test BS, generated from trained HSMM and HMM. HSMM and HMM were both first trained on the same dataset simulated using the uniform duration distribution in red (mean 50 a.u. and width 6 a.u.). Subsequently sequences of BSs were generated from each trained model. The geometric distribution implicit in HMM, produced BS durations across the whole range of possible values (green bars). The explicit normal duration distribution in HSMM concentrated the durations in the correct range of values (blue bars). The best fitted geometric (green line) and normal (blue line) BS duration distributions respectively for the HMM and the HSMM, are also shown.

Plots of the JSD between the histogram of durations generated by each model and the histogram of durations underlying the training data for the test BS, are depicted in Figure 5 as a function of the BS mean duration at two different SNR of the training dataset. HSMM was consistently more accurate and more robust at reproducing the true duration distributions, as evidenced by the lower JSD values across all mean duration and SNR values. Notably, HMM reproducibility deteriorated as mean duration increased. This suggests that HMM estimation of duration was more sensitive to the amount of data available to infer BS transitions accurately. That is, given a fixed length of data, a longer BS duration translates into a lower number of state transitions. In this situation, the ability of HSMM to capture the actual dynamics of the underlying system helps to regularise and stabilise the inference. No apparent effect of the SNR of the data was found for any of the two models.

**Figure 5.**
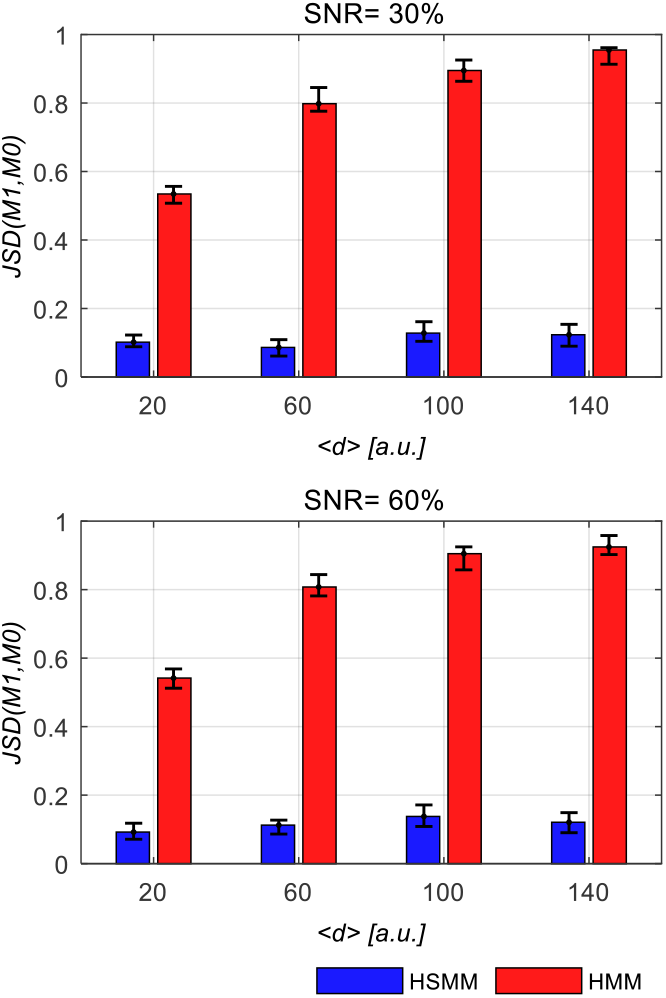
Jensen-Shannon Divergence (JSD) between training and model generated BS durations in HSMM and HMM. The mean duration of a test BS was varied at two SNR values. A lower JSD value indicates closer resemblance between the duration distribution generated by the model and the duration distribution underlying the training dataset. HSMM consistently and more robustly regenerates the BS duration distribution of the true underlying system. The error bars over 100 realisations of the experiment are indicated.

The above results demonstrate that, although the standard HMM can accurately fit the data produced by a non-Markovian HDS, it fails to capture the actual dynamics of the hidden BS process. Therefore, when prediction and interpretation is the goal, using a HMM to describe a non-Markovian system (as seems to be the case of the brain) can lead to wrong conclusions. In the next sections we will evaluate the performance of HSMM vs HMM for the analysis of a non-Markovian HDS, in situations where accurate estimation of the underlying BS duration model is important.

### 3.2 BS allocation under increased data uncertainty

HSMM and HMM were compared in terms of their accuracy in identifying the hidden BS sequence underlying a previously “unseen” (out-of-sample) EEG test dataset. This amounts to solving a decoding problem. That is, given a previously trained model HSMM (or HMM), with estimated parameters 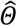, and a new observation sequence *y*_1:*T*_ (test dataset), what is the most probable BS path that produced the observations? Such a task is solved using the well-known Viterbi algorithm (Yu, 2015). In this simulation experiment, the test dataset was noisier than the training dataset. This simulation mimics a situation in which a previously trained model is used to monitor BSs under uncontrolled (unexpected) fluctuations of the observation noise level. In such situation, one wished the model was robust to the noise fluctuations. One hundred pairs of training and test datasets were simulated from a HDS with 3 BSs (see section 2.4), for each SNR value of the training and the test datasets. Durations of all BSs were sampled from a uniform distribution of width 6 a.u. Apart from the SNR, the parameters of the underlying HDS generating the training and the test datasets in each pair were identical. As in the previous simulations, different memory lengths of the underlying BS sequence were explored by varying the mean duration of a test BS. The mean duration of the remaining BSs was fix at 10 a.u.

Figure 6 shows plots of the SAS for HSMM and HMM as a function of the mean duration of the test BS, and for different SNR values of the training and the test datasets. For increased difference between the SNR of the training and the test datasets, the accuracy of the HMM deteriorated significantly, while HSMM remained stable with a SAS above 90% in almost all cases. Interestingly, in the case of greatest reduction in data quality (Figure 6 lower panel) the accuracy of the HSMM increased, while the HMM deteriorated for increasing values of the mean BS duration.

**Figure 6.**
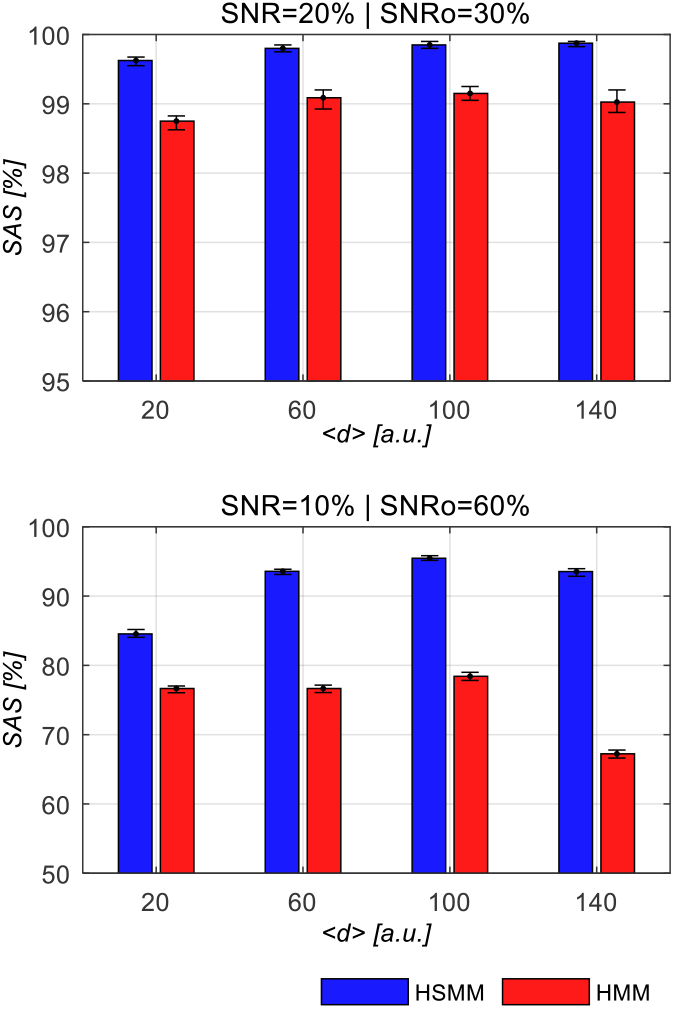
Accuracy of BS allocation based on unreliable (SNR reduced) data. HSMM and HMM were first trained on data produced by a non-Markovian system and were then used to decode the BS sequence underlying a new test dataset from the same system but with reduced SNR. The mean duration of a test BS was varied at two levels of SNR of the training dataset (SNR_0_) and the test dataset (SNR). The accuracy of the decoding was evaluated using the Sequence Accuracy Score (SAS), between the decoded and the true BS sequence. Higher SAS indicates a more accurate decoding. HSMM achieved higher SAS for all duration values explored. The error bars over 100 realisations of the experiment are indicated.

### 3.3 Allocation of *Cognitive States* (CS)

We have defined a BS as a mode of operation (i.e. a pattern of neural activity or functional network) that remains stable for a short period of time (e.g. below 500ms (Baker et al., 2014)) before rapidly transitioning (switching) to a different mode. On the other hand, longer lasting, more complex mental or cognitive states (CS) could be seen as a specific sequence of these short-lived BSs. Therefore, identifying data segments corresponding to BS sequences with distinctive dynamical structures is paramount for the detection and monitoring of such CS. This task amounts to solving a model evaluation problem (Bishop, 2006). That is, given a trained model HSMM (or HMM) with estimated parameters 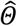, and given a sequence of observations *y*_1:*T*_, we are interested in the probability 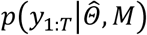 that the observations were generated by the model. This is the conditional probability of the data after marginalisation over all possible BS paths, which can be used to compute the discriminative power between two models (see section 2.5 and Appendix B.5).

HSMM and HMM were compared in terms of their ability to identify a target CS among a collection of candidate CSs. The candidate CSs differed in the mean duration of the component BSs. This experiment mimics a situation in which we would like to for example discriminate between a healthy population and a clinical group, where the clinical group is characterised by an alteration in the time the brain spends in each BS. Labelled data were first used to train different models, where each label denoted a specific CS so that each trained model (HSMM or HMM) was associated with a unique CS. A new unlabelled data segment was then allocated to a specific CS by computing the probability of each model having generated the data, and assigning the label associated with the model of highest probability.

The labelled training dataset of each CS consisted of a data sequence, generated from a non-Markovian HDS with 3 BSs. Different CSs were obtained by varying the mean duration of the 3 BSs from 2 a.u. to 22 a.u. (step-size 2). The unlabelled dataset was generated in the same way with mean BS duration of 10 a.u. for all the states. In all simulated data sequences the width of the BS duration distributions were all fixed to 6 a.u. The training and allocation procedure described above was then performed for HSMM and HMM separately on the same datasets and the whole experiment was repeated 100 times. Again, the two models achieved an excellent goodness of fit in all training sessions (results not shown).

The ability of the models to allocate the test CS correctly was measured in terms of the DPS (section 2.5). Figure 7 shows the *DPS* as a function of the mean BS duration encoding each of the candidate CSs, for both HSMM and HMM. Each point of the curve represents the average information per test data sample to discriminate in favour of the hypothesis *H*_0_ that the test CS was generated by the candidate model tested, against the hypothesis *H*_1_ that it was generated by the correct model (BS duration 10 a.u.). The zero value represents no ability to discriminate between the two hypotheses and therefore indicates the CS the test sequence is allocated to. Discrimination based on HSMM allowed for correctly allocating the test sequence to the CS associated with the model with mean BS durations of 10 a.u. In the case of HMM, there was no information in the test data that allowed discriminating the correct CS from any of the candidate ones, which made CS allocation not possible.

**Figure 7.**
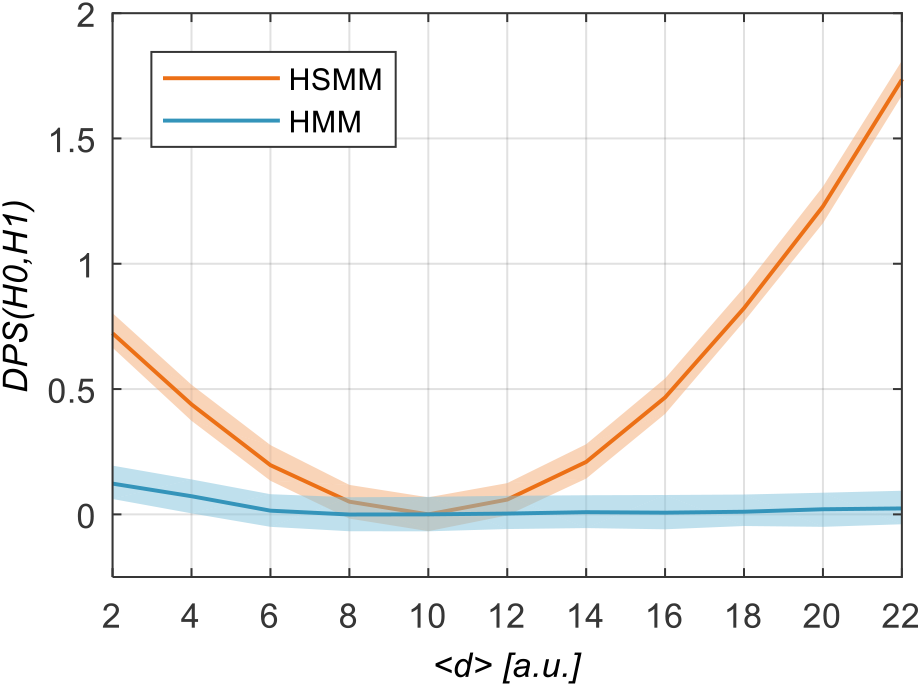
Discriminative Power of HSMM vs HMM to allocate data segments. HSMM vs HMM were trained on data segments associated with BS sequences (Cognitive States (CS)) that differed in the mean duration of the constituent BSs. A new unlabelled data segment (a CS with mean BSs duration of 10 a.u.) was then decoded with each of the trained models and allocated to the CS associated with the model with highest probability of generating the unlabelled data. Each point on the curves is the Discriminative Power Score, DPS, a measure of the discrimination in favour of the hypothesis H0 that the test data was generated by a candidate model (with duration on the x-axis), against the hypothesis H1 that it was generated by its true model (〈*d*〉 = 10). A value of *DPS* = 0 indicates no discrimination between H0 and H1, that is the candidate model is very close to the truth. The shadowed areas represent the 95% confidence interval for discrimination. HSMM is able to allocate the test sequence to the correct CS, whereas HMM has no discrimination power.

### 3.4 Resting-state EEG: light-tail vs heavy-tail duration distributions

We have so far shown that the main advantage of HSMM over HMM is that it allows for the explicit modelling of BS durations. Here we use the real resting state EEG data to compare between alternative duration models. Specifically, we compare using HSMM with Log-Normal (heavy-tailed and skewed) vs Normal (light-tailed and symmetric) BS duration distributions to model the fluctuations of the EEG amplitude’s envelopes.

First we investigate whether there is evidence in the data supporting one duration model vs the other. Two HSMMs, each using one of the duration models with maximum duration *D* = 5000*ms*, were trained on the actual data as described in section 2.6. In the two cases, 4 BSs were automatically inferred from data using model comparison. Figure 8 shows the convergence curve of the NFE for the two optimal models with 4 BSs. The HSMM with Log-Normal durations converged faster and showed higher model quality with a log-Bayes Factor of 400 in its favour after convergence.

**Figure 8.**
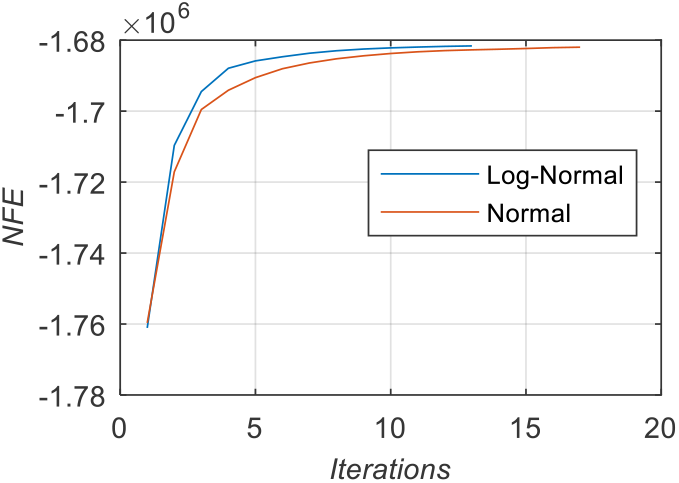
Convergence of the VB algorithm. Negative Free Energy (*NFE*) of the HSMM for the two duration models used in the EEG resting-state data. The HSMM with Log-Normal durations converged faster and reached a higher *NFE* value.

The partial correlation maps associated with each BS are shown in Figure 9. Based on the pair-wise similarity between the maps of the two duration models, the BSs were relabelled and matched based on these maps by simple visual inspection. The maps from BS1 and BS2 are approximately spatially homogeneous with opposite signs, while maps from BS3 and BS4 show more spatial features. Resting-state EEG is mainly dominated by global alpha activity, which has been characterised as a bi-stable system jumping erratically between low-power and high-power modes (Roberts et al., 2015). Maps from BS1 and BS2 could then be reflecting the dominant alpha fluctuations, with BS1 and BS2 representing the low- and high-power alpha modes respectively. BS3 and BS4 can then be related to more local “bursting” activity in lower and higher frequency bands.

**Figure 9.**
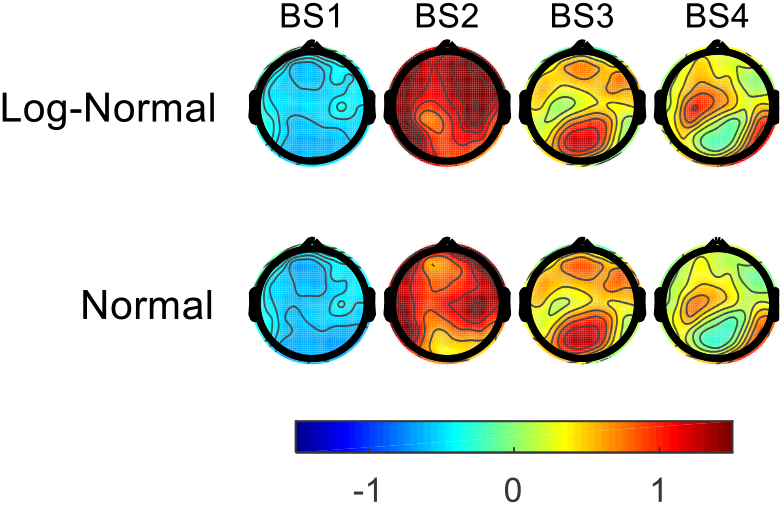
BSs allocation in resting-state EEG. Partial correlation maps associated with the estimated BSs. The duration model used to obtain each row is also indicated.

Figure 10 summarises the temporal characteristics of the inferred BSs in terms of their duration distributions induced by the posterior expected values of the parameters of the Log-Normal and the Normal duration models; as well as the corresponding fractional occupancy of the BSs. The duration distributions of the two models showed consistent behaviour. BSs with heavier-tailed Log-Normal distributions also show more right-shifted and wider Normal distributions. Notably, the standard deviation of the distribution of each BS, scales with its mean, with BS1 and BS2 showing the most striking differences. For completeness, several statistics of the wining Log-Normal duration model for the 4 BSs are reported in Table 1.

**Figure 10.**
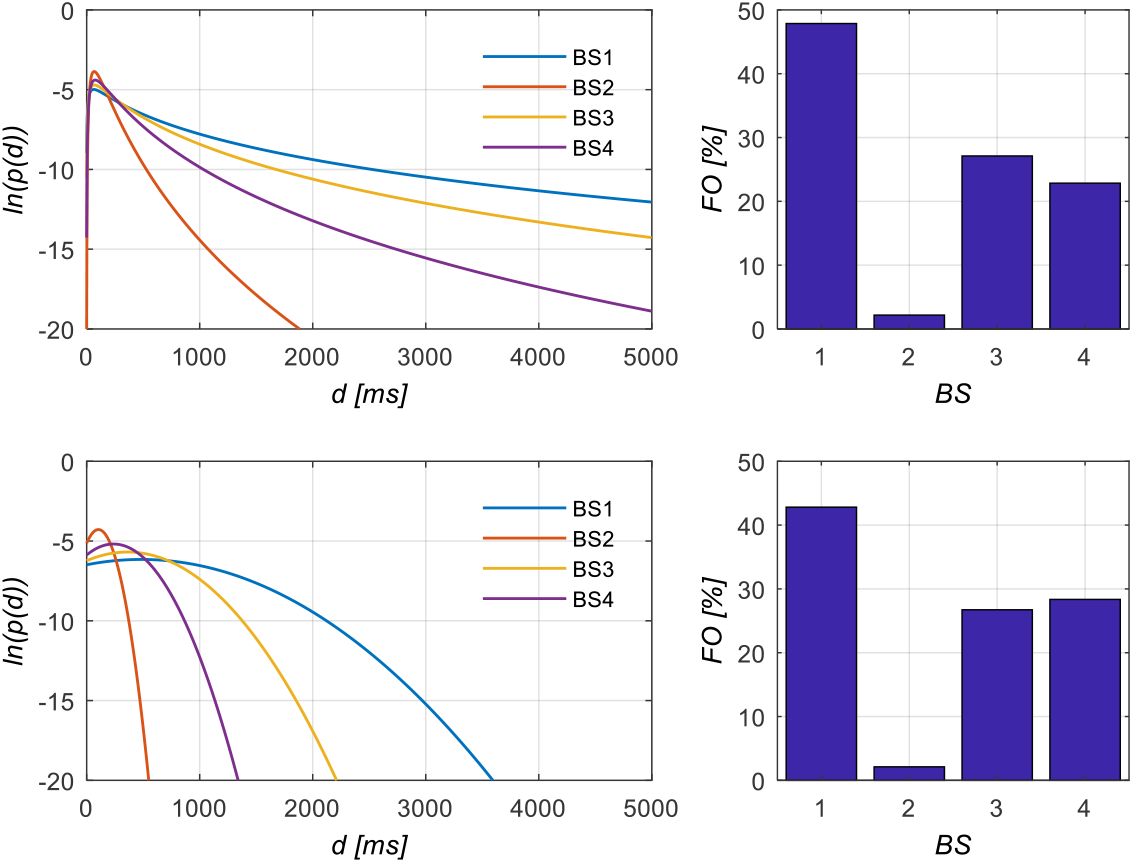
Logarithm of duration distributions (*In*(*p*(*d*))) and Fractional Occupancy (*FO*) of BSs in resting-state EEG. Log-Normal (top) and Normal (bottom) duration distributions induced by the posterior estimates of the parameters of each model; and the corresponding fractional occupancy of the BSs. The duration axis has been truncated at 1400ms to visualise the effects of interest.

**Table 1.** Statistics of the Log-Normal distribution of durations of all BSs. STD, SKW, eKRT and CV denote standard deviation, skewness, excess Kurtosis and coefficient of variation, respectively. Means, medians, modes and STDs are expressed in milliseconds.

| BS | Mean | Median | Mode | STD | SKW | eKRT | CV |
| --- | --- | --- | --- | --- | --- | --- | --- |
| 1 | 448.9 | 246.1 | 65.3 | 586.6 | 9.7 | 352.8 | 1.30 |
| 2 | 114.7 | 96.8 | 69.0 | 72.9 | 2.16 | 9.3 | 0.64 |
| 3 | 290.4 | 181.8 | 70.3 | 347.5 | 5.77 | 93.3 | 1.20 |
| 4 | 190.7 | 141.4 | 77.7 | 172.5 | 3.5 | 26.9 | 0.90 |

The FO shows regularities that are partially supported by the duration distributions. Overall, BSs with heavier Log-Normal tail and greater Normal right-shift also show higher FO. This trend might be expected because a duration distribution with heavier tail (greater right-shift) allocates higher probability mass at longer durations, which should correspond to higher FO. The exception is the FO corresponding to the Normal model of BS3 and BS4, for which smaller right-shift of BS4 (as compared to BS3) translates into a higher FO.

The relationship between tail heaviness and FO does not necessarily hold in general because the FO index conflates two different aspects of BS dynamics: state durations and transition probabilities. This can explain the apparent BS3/BS4 anomaly. The transition probabilities in Figure 11, show that in the case of the Normal duration model, the probability of entering BS4 is increased as compared to the log-normal case, while the probability of entering BS3 is reduced.

**Figure 11.**
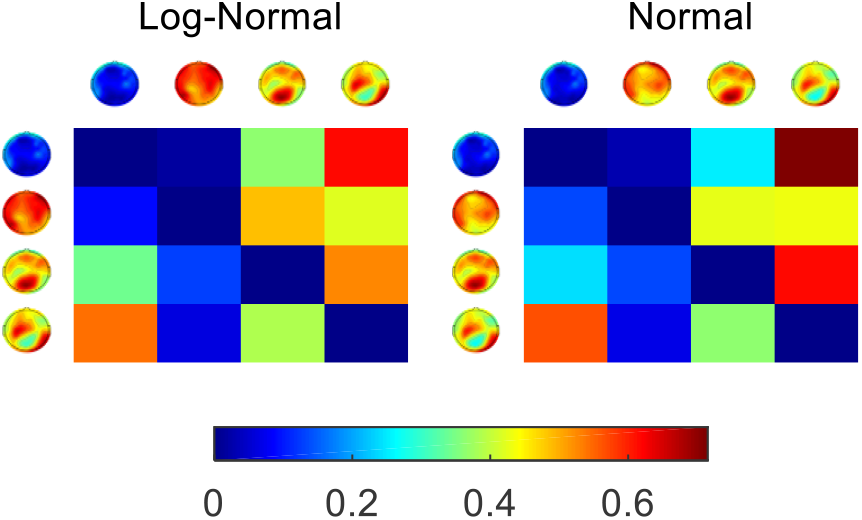
Transitions between BSs in resting-state EEG. Posterior estimates of the transition matrices of the HSMMs for the two duration models.

The above rationale is made explicit in Figure 12 showing the stationary (equilibrium) probability distribution of the Markov chain embedded in the HSMM (super-state representation). These probabilities can be interpreted as the long-run proportion of visits to each BS per unit time. In the case of the Normal duration model the proportion of visits to BS4 is greater than the visits to BS3, while the reverse holds for the Log-Normal model. Note however that the FO trends cannot be completely explained by the stationary distribution of BSs alone either. That is, a greater visiting rate to a BS does not always correspond to greater FO (e.g. BS1 vs BS4 in Figure 10 and Figure 12).

**Figure 12.**
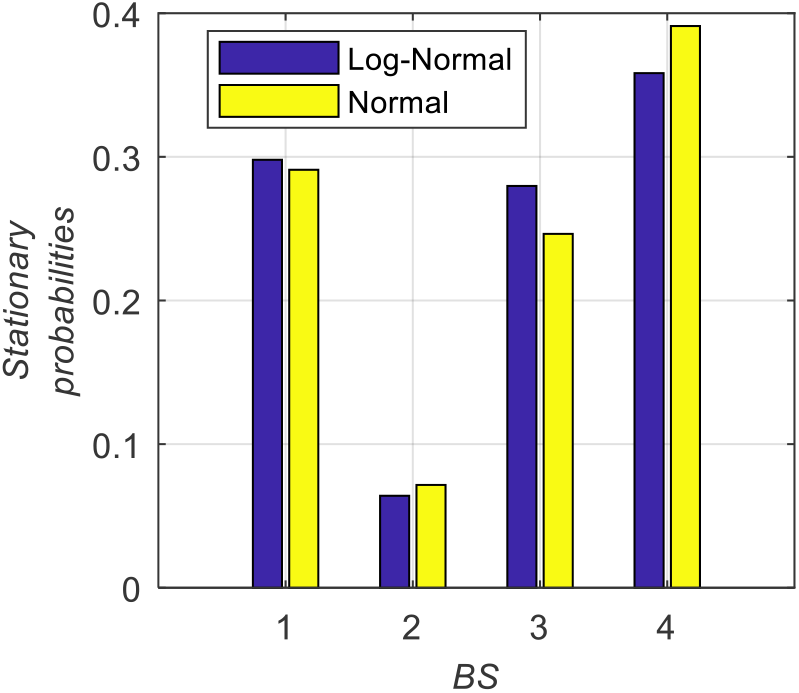
Stationary (equilibrium) distribution of the Markov chain embedded in HSMM. These probabilities highlight the differences between the BSs in terms of the frequency with which they are visited.

## 4 Discussion

We have proposed Hidden Semi-Markov Model (HSMM) as a generative model for Brain State (BS) allocation in M/EEG, which allows for the explicit modelling of the BS duration. The added flexibility circumvents a well-known limitation of the standard HMM where the implicit distribution of BS duration is Geometric. The Geometric distribution places more probability mass at shorter than at longer BS duration, making HMM intrinsically biased towards fast BS switching. A more fundamental problem is that the Geometric distribution is *memoryless.* This property implies that the probability of waiting time until the next BS transition is independent of the time already spent in the current BS. For a model to describe such behaviour, it must continuously *forget* the state the system is currently in (hence memoryless). A Geometric duration is then inconsistent with the long-range temporal dependency (“long memory effect”) and near scale-invariance (self-similarity) ubiquitous in real EEG data (Buzsáki and Mizuseki, 2014; Freeman, 2009; Kello et al., 2010; Linkenkaer-Hansen et al., 2001; Roberts et al., 2015). This poses strong limitations on using HMM for modelling this data accurately.

The ability of HSMM to produce sensible non-Markovian behaviour was demonstrated by generating synthetic BS sequences from an HSMM with Uniform BS duration PDFs. The simulated sequences showed non-Markovian temporal dependencies previously observed in empirical microstates sequences (von Wegner et al., 2017) (Figure 3a & Figure 3c). The observed temporal dependencies were sensitive to variations of the two BS duration parameters (mean and width of the PDF), indicating that these parameters are identifiable from the data. In contrast, the lack of sensitivity of HMM to changes in the duration parameters (Figure 3b & Figure 3d) suggested that HMM cannot capture the temporal features of BS in data with non-Markovian behaviour. The data simulation test (Figure 4 and Figure 5) demonstrated this hypothesis by showing that, although HSMM and HMM achieved comparable goodness-of-fit during model training on data with non-Markovian properties, only HSMM was capable of re-generating BS duration PDFs comparable to those of the original non-Markovian system.

Another usually overlooked limitation of HMM is that it places a small penalty on adding a new state, and no penalty on the new state having similar features to another state, nor on the state sequence rapidly switching between the new and a similar existing state (Fox et al., 2011). Consequently, using model comparison to infer the number of states based on data with non-Markovian behaviour, can lead to the creation of unnecessary extra states and unrealistically fast BS switching, in order to reproduce long states. Indeed, previous works using HMM for BS allocation have found the Negative Free Energy to increase monotonically with the number of BSs (Baker et al., 2014; Rukat et al., 2016). The fast-switching can also harm the reliable estimation of the model, since parameters are informed by fewer data points. This means that stable inference using HMM requires large amount of data, making single subject inference and random effect analysis, challenging. In practice, BS allocation with HMM is then typically applied to group data, where a single HMM is fitted to the whole (subject-concatenated) dataset (Vidaurre et al., 2017). In the present paper we obtained meaningful and reliable results using HSMM to analyse resting-state data from a single subject. Importantly, the number of BSs was automatically estimated from the data via NFE maximisation. Note that the redundant BSs and fast switching in HMM is not necessarily a problem if model averaging is the goal (Woolrich et al., 2013). However, if interpretation is important, one would like to incorporate prior knowledge that a “slow” switching dynamics is more likely. HMM does not incorporate this kind of prior, while in HSMM this is done explicitly.

To further characterise HSMM and HMM, we compared their performance for BS allocation in two tasks where accurate prediction and interpretation was important. The first task tested whether an accurate modelling of BS durations aided BS allocation based on data with increased external noise. The (Bayesian) rationale is that if the data is unreliable, the accuracy and robustness of the inference (here BS decoding) is determined by the ability of the model to represent the actual system producing the observations, so that noise is effectively separated from the system dynamics. We found that HSMM allowed robust and accurate allocation of BSs for different data uncertainty levels and different mean BS durations (Figure 6). This is because the Normal duration used in HSMM was a good approximation of the true BS duration PDF (Figure 4). In contrast, HMM was not accurate enough to overcome the effect of the increased data uncertainty, which led to non-robust inference dominated by the noise fluctuations.

The second task involved identifying characteristic BS sequences containing information not reducible to the individual contribution of the component BS. We have called these sequences Cognitive States (CS), indicating that it might be a more appropriate representation of a “mind state”, than an individual short-lived BS. Concepts similar to the CS has been used to interpret both task-related (Borst and Anderson, 2015; Laganaro, 2017) and resting-state (Michel and Koenig, 2018; Vidaurre et al., 2017) activity. The CS to be identified differed from the candidate CSs only in the mean duration of their component BSs. That is, is the implicit Geometric distribution in HMM enough to discriminate between CSs, or a more accurate representation of BS duration is required? Our results showed that, while HSMM allowed correct and robust allocation of an unlabelled CS, HMM could not discriminate between the unlabelled CS and any of the candidate CSs. Intuitively it can be argued that in the identification of long-lived CSs, the temporal dependencies might play an even more important role than in the identification of individual short-lived BS, and therefore HSMM is recommended over HMM.

We have demonstrated that accurate modelling of durations is paramount for BS allocation. So which duration distribution should we use? The choice of duration PDF determines the temporal properties of the BS process, which will ultimately impose constraints on the kind of dynamics that the model can explain. A light-tailed “bell-shaped” PDF is typical of a process with a characteristic time scale (e.g. periodicity), while a heavy-tailed right skewed PDF corresponds to a process showing long-term dependencies and temporal scaling law (Gschwind et al., 2015; Linkenkaer-Hansen et al., 2001; Roberts et al., 2015). Importantly, these temporal features are indicative of specific non-linear dynamical mechanisms generating the data (see (Cocchi et al., 2017) for a review). We can therefore use the explicit BS duration PDF in HSMM to express alternative hypotheses of temporal dependency and scaling-law of the BS sequence, and select the best distribution given the EEG data in a principled way via Bayesian Model Selection. This approach can bridge between the observed BS data features and the associated underlying neural mechanisms, which would allow interrogating the data about such mechanisms. To illustrate the approach we used actual resting-state EEG data to select between a heavy-tailed (Log-Normal) and a light-tailed (Normal) BS duration PDF for BS allocation using HSMM. Model comparison demonstrated conclusive evidence in favour of the Log-Normal PDF. A closer analysis revealed that two of the states (BS1 and BS3) showed signs of scaling-law dynamics, while the other two (BS2 and BS4) showed a sharper peak more consistent with a characteristic time scale (Figure 10). This suggests that a mixture of heavy-tailed and light-tailed distributions might be a duration model worth exploring in the future. Favouring a heavy-tailed model also indicates that large duration values are observed more often than expected from Gaussian statistics (see Table 1). The relatively high mean durations reported here should then be contrasted with those obtained using HMM to analyse envelope fluctuations of the wide band EEG (Hunyadi et al., 2019; Rukat et al., 2016)

Finally, we showed that, in the context of HSMM, post-hoc summary statistics such as FO (Baker et al., 2014; Vidaurre et al., 2018, 2017), which conflate BS duration and transition probabilities, are difficult to interpret because they reflect redundant and conflicting information. In HSMM, observed changes in FO are disambiguated because durations and BS transitions are modelled explicitly and independent of each other, so that their individual effects can be assessed. This implies that experimental or clinical conditions associated with changes in BS durations, can be separated from those associated with altered transition probabilities between BSs. In HMM this is not possible because BS transitions and durations are also confounded. Our results in actual data show that, in the case of the Log-Normal durations, the FO reflected the heaviness of the tail of the PDF, so that total occupancy was dominated by the dwell time in each BS visit, rather than by the frequency of the visits. In the case of Normal durations, FO seems to reflect a combination of the two factors.

## 5 Acknowledgments

We thank Aland Astudillo for providing assistance in the pre-processing of the experimental data presented. N.J.T-B. is funded by the Engineering and Physical Sciences Research Council (EPSRC) Grant EP/N006771/1. D.A. is funded by a PhD scholarship from CONICYT, Chile 21170611. W.E-D. is funded by CONICYT, Chile, Projects FONDECYT 1161378, ANILLO ACT172121 and Basal FB0008. N.J.T-B and W.E-D contributed equally to this research.

## Appendix A Definition of Signal-to-Noise Ratio

For a given data sequence, the SNR was defined as the percentage of noise-free signal energy present in the data:

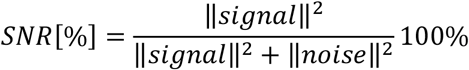

For iid Gaussian noise in each channel and normalised BS maps, it can be demonstrated that

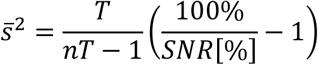

Where 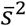 is the unbiased sample variance estimate and *n* and *T* are the number of channels and the length of the data sequence respectively. This expression was then used to compute the simulated noise variance required to achieve a selected SNR value.

## Appendix B Evaluation scores

### B.1 Auto Information Function

The Auto Information Function (AIF) is analogous to the autocorrelation function, but for symbolic sequences such as the BSs sequences. It measures the dependence between different time points with a given time lag *l*. This is done by measuring the Kullback-Leibler divergence between the symbol distributions at time *t* and *t* + *l* That is, the AIF for time lag *l* is defined as

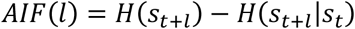

This is the difference between the entropies *H* of the distributions *p*(*s_t+l_*) and *p*(*s_t+l_*|*s_t_*). For a stationary Markov chain the *AIF* can be obtained analytically (von Wegner et al., 2018).

### B.2 Sequence Accuracy Score

Denoting the BS decoded at time *t* as *ŝ_t_* and the true one as 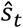, the Sequence Accuracy Score (SAS) between the sequences *ŝ*_1:*T*_ and 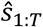 of equal length *T* is defined as

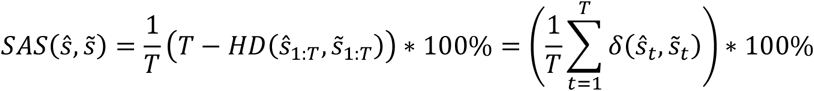

where *δ*(·,·) is the Kronecker’s delta function and *HD*(*x,y*) denotes the Hamming distance between sequences *x* and *y* (Navarro, 2001).

### B.3 Jensen-Shannon Divergence

In probability theory, the Jansen-Shannon Divergence (JSD) (Endres and Schindelin, 2003) is used to measure the similarity or dissimilarity between two probability distributions or histograms. JSD is a symmetrised version of the Kullback-Leibler divergence *KL*(*a*||*b*) between two distributions *p* and *q*. It is defined as

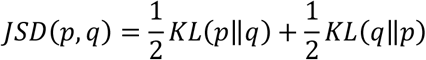

The *JSD* is bounded between 0 (perfect similarity) and 1 (perfect dissimilarity) and its square root is a metric known as the Jensen-Shannon distance..

### B.4 Log-Bayes Factor)

Given a data sequence *y*_1:*T*_ and two given model classes *M*_1_ and *M*_0_, the log-Bayes factor (Kass and Raftery, 1995) is defined as

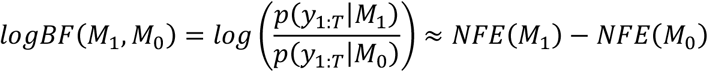

where *p*(*y*_1:*T*_|*M_k_*) is the evidence of model class *M_k_*. Since the evidence cannot be calculated in closed form, we used its lower bound approximation in terms of the (variational) Negative Free Energy, *p*(*y*_1:*T*_|*M_k_*)≈ *NFE*(*M_k_*).

### B.5 Discriminative Power Score

Given a data sequence *y*_1:*T*_, and two models (of the same model class) each defined by the set of parameters *Θ*_1_ or *Θ*_0_, the ease of discriminating between the two models is defined in terms of the probabilistic distance (Juang and Rabiner, 1985)

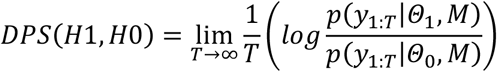

where *p*(*y*_1:*T*_|*Θ_k_,M*) is the probability that the data sequence was generated by the model *k* (i.e. the model with parameters *Θ_k_*), after marginalization over all possible BS paths. That is, for a sufficiently long sequence, if *H*1 and *H*0 are the hypotheses that the data sequence was generated by model 0 and model 1, respectively, *DPS*(*H*_1_, *H*_0_) is the average information per data sample for discrimination in favour of *H*_1_ against *H*_0_.

## Notes

https://github.com/daraya78/BSD

