## Supplementary material for "The discrete logic of the Brain - Explicit modelling of Brain State durations in EEG and MEG": Suplemental Methods

### Supplements

#### 1 Mean Field Approximation and Negative Free Energy

The mathematics of the VB framework for HSMM is described in detail in (Hudson, 2009). The mean-field approximation used here has the following form:

$$q(s_{1:T}, r_{1:T}, \pi, A, \theta, \lambda) = q(s_{1:T}, r_{1:T}) q(\pi) \left[ \prod_{k=1}^{K} q(a_{k:}) \right] q(\mu) q(\Sigma) q(\tau) q(\rho)$$

where, in addition to the factorisation between hidden states and parameters (see main text) we have assumed factorisation over the parameters of both the duration and the emission distributions; as well as over the rows of the transition matrix (Beal, 2003). This factorisation induces the following form of the NFE:

$$\begin{aligned} NFE\big(q(s_{1:T}, \Theta)\big) = -KL(q(\pi)\|p(\pi)) - \sum_{k=1}^{K} KL(q(a_{k:})\|p(a_{k:})) - \\ - \sum_{k=1}^{K} KL\big(q(\mu^{(k)})\|p(\mu^{(k)})\big) - \sum_{k=1}^{K} KL\big(q(\Sigma^{(k)})\|p(\Sigma^{(k)})\big) - \\ - \sum_{k=1}^{K} KL\big(q(\tau^{(k)})\|p(\tau^{(k)})\big) - \sum_{k=1}^{K} KL\big(q(\rho^{(k)})\|p(\rho^{(k)})\big) + \\ + logC(s_{1:T}) \end{aligned}$$

where $KL(q(x)\|p(x))$ denotes the Kullback-Leibler divergence between the approximate variational posterior $q(x)$ and the prior $p(x)$; and $C(s_{1:T})$ is a normalisation constant that can be computed after each VB-E step (Hudson, 2009). Note that due to conjugacy of the priors, all KL divergences are between known standard distributions and can be computed analytically. We refrain from including the KL expressions here, since these are standard results available somewhere else.

#### 2 Priors and posterior updates for all the parameters

This section presents the update equations of the VB-M-Step. It uses the following marginal statistics obtained from the Forward-Backward algorithm implemented in the VB-E step:

- The joint posterior probability of the joint process $(s_t, r_t)$ being at $(i, d)$ at time $t-1$ and at $(j, d')$ at time $t$:

$$\xi_t(i, d, j, d') = q\big((s_{t-1}, r_{t-1}) = (i, d), (s_t, r_t) = (j, d')\big)$$

- The posterior probability that the joint process $(s_t, r_t)$ being at $(i, d)$ at time $t$

$$\gamma_t(i, d) = q\big((s_t, r_t) = (i, d)\big) = \sum_{d'=1}^{D} \sum_{j=1}^{K} \xi_t(j, d', i, d)$$

Given the structure of the HSMM presented here, these statistics can usually be reduced respectively to $\xi_t(i, 1, j, d)$, the posterior probability of a transition occurring at time $t$; and $\gamma_t(i) = q(s_t = i) = \sum_{d=1}^{D} \gamma_t(i, d)$, the posterior probability of state $i$ being active at time $t$.

#### 2.1 Multivariate Normal emission models

The signal emitted from the $k$-th BS is modelled as a $n$-variate Normal distribution with mean $\mu^{(k)}$ and precision matrix $\Sigma^{(k)}$.

$$p\left(y_t \middle| s_t = k, \mu^{(1:K)}, \Sigma^{(1:K)}\right) = N_n\left(y_t; \mu^{(k)}, {\Sigma^{(k)}}^{-1}\right)$$

For the simulation experiments, the data was iid with BS precision matrix $\Sigma^{(k)} = diag\left(\sigma^{(k)}\right)$, where $\sigma^{(k)}$ is the $n \times 1$ vector of precisions (inverse variances) of each channel's data. For the actual data a full precision matrix was assumed.

##### 2.1.1 Prior distribution for means of emissions

For both the simulations and the actual data we assumed that all BS means are independent across states and iid a priori with identical conjugate weakly informative priors

$$p(\mu|\mu_0, \Sigma_0) = \prod_{k=1}^{K} N_n\left(\mu^{(k)}; \mu_0, \Sigma_0^{-1}\right), \quad \mu_0 = 0_{n\times 1}, \quad \Sigma_0 = \sigma_0 I_n$$

Where $0_{n\times 1}$ is a $n \times 1$ vector of zeros, $I_n$ is the identity matrix of size $n \times n$. We used $\sigma_0 = 0.1$ for the simulations and $\sigma_0 = 0.01$ for the actual data.

##### 2.1.2 Prior distribution for precision of emissions

For the iid emission model used in simulations, precisions were assumed to be are independent over states and channels a priori with identical conjugate Gamma priors, and weakly informative shape and scale parameters $a$ and $b$, respectively

$$p(\sigma|a_0, b_0) = \prod_{k=1}^{K} \prod_{i=1}^{n} Gam\left(\sigma_i^{(k)}; a_0, b_0\right)$$

where $a_0 = 0.001$ and $b_0 = 1000$. For the full precision model used in the actual data, precision matrices are independent over states a priori with identical conjugate $n$-dimensional Wishart distributions with degrees of freedom $a$ and scale matrix $B$

$$p(\Sigma|a, B) = \prod_{k=1}^{K} W_n(\Sigma^{(k)}; a_0, B_0)$$

Where $a_0 = n$ and $B_0 = nI_n$.

##### 2.1.3 Posterior updates for means of emissions

Due to conjugacy of the priors, the variational densities of the states' means of the two emission models have both Gaussian forms. For the iid model the posterior is also iid

$$q(\mu) = \prod_{k=1}^{K} N_n\left(\mu^{(k)}; \hat{\mu}^{(k)}, \left(diag\left(\hat{\sigma}^{(k)}\right)\right)^{-1}\right)$$

With posterior mean and precisions given by

$$\hat{\mu}_i^{(k)} = \left(\sigma_0 \mu_{0_i} + \hat{a}_i^{(k)} \hat{b}_i^{(k)} \sum_{t=1}^{T} \gamma_t(k) y_{t_i}\right) \left(\sigma_0 + N^{(k)} \hat{a}_i^{(k)} \hat{b}_i^{(k)}\right)^{-1}, \qquad \begin{matrix} i = 1, \dots, n \\ k = 1, \dots, K \end{matrix}$$

$$\hat{\sigma}_i^{(k)} = \sigma_0 + N^{(k)} \hat{a}_i^{(k)} \hat{b}_i^{(k)}$$

Where $\hat{a}_i^{(k)}$ and $\hat{b}_i^{(k)}$ are the shape and scale parameters of the approximate posterior of the precisions, associated with emissions from state $k$ (see below) at channel $i$; $N^{(k)}$ are the expected state's counts (i.e. the expected number of states' visits)

$$N^{(k)} := \sum_{t=1}^{T} \gamma_t(k), \qquad k = 1, \dots, K$$

In the case of the emission model with full precisions, the variational posterior for the means are independent over states so that

$$q(\mu) = \prod_{k=1}^{K} q\left(\mu^{(k)}\right) = \prod_{k=1}^{K} N_n\left(\mu^{(k)}; \hat{\mu}^{(k)}, \hat{\Sigma}^{(k)^{-1}}\right)$$

With posterior parameters

$$\hat{\mu}^{(k)} = \left(\Sigma_0 \mu_0 + \hat{a}^{(k)} \hat{B}^{(k)} \sum_{t=1}^{T} \gamma_t(k) y_t\right) \left(\Sigma_0 + N^{(k)} \hat{a}^{(k)} \hat{B}^{(k)}\right)^{-1}, \qquad k = 1, \dots, K$$

$$\hat{\Sigma}^{(K)} = \Sigma_0 + N^{(k)} \hat{a}^{(k)} \hat{B}^{(k)}$$

##### 2.1.4 Posterior updates for precisions of emissions

Again, conjugacy of the priors ensures that the variational posteriors have the same functional form as the prior. In the case of the iid observation model, the variational posterior of the precisions factorises over states and channels so that

$$q(\sigma) = \prod_{k=1}^{K} \prod_{i=1}^{n} q\left(\sigma_i^{(k)}\right) = \prod_{k=1}^{K} \prod_{i=1}^{n} Gam\left(\sigma_i^{(k)}; \hat{a}_i^{(k)}, \hat{b}_i^{(k)}\right)$$

With posterior shape and scale parameters

$$\hat{a}_i^{(k)} = a_0 + \frac{1}{2} N^{(k)}$$

$$\hat{b}_i^{(k)} = \left(b_0^{-1} + \sum_{t=1}^{T} \left(\gamma_t(k) \left(y_{t_j} - \hat{\mu}_i^{(k)}\right)^2 + \hat{\sigma}_i^{(k)^{-1}}\right)\right)^{-1} \qquad \begin{matrix} i = 1, \dots, n \\ k = 1, \dots, K \end{matrix}$$

In the case of the emission model with full precision matrices, the variational posterior of the precisions factorises over states with state-specific Wishart distributions so that

$$q(\Sigma) = \prod_{k=1}^{K} q\left(\Sigma^{(k)}\right) = \prod_{k=1}^{K} W_n\left(\Sigma^{(k)}, \hat{a}^{(k)}, \hat{B}^{(k)}\right)$$

With posterior degrees of freedom and scale parameters

$$\hat{a}^{(k)} = a_0 + N^{(k)}$$
$$\hat{B}^{(k)} = \left( B_0^{-1} + \sum_{t=1}^{T} \gamma_t(k) \left(y_t - \hat{\mu}^{(k)}\right)^T \left(y_t - \hat{\mu}^{(k)}\right) + tr\left(\hat{\Sigma}^{(k)^{-1}}\right) \right)^{-1}, \qquad \forall k$$

#### 2.2 Duration models

##### 2.2.1 Normal duration model

We impose independent conjugate and weakly informative prior distributions on the mean and the precision associated with each state.

###### *2.2.1.1 Prior distribution for the parameters*

Independent Normal prior distributions with identical mean $\tau_0$ and precision $\delta_0$ were assumed on the mean duration of each state so that the full prior factorises over states:

$$p(\tau) = \prod_{k=1}^{K} p\left(\tau^{(k)} \middle| \tau_0, \eta_0^{-1}\right) = \prod_{k=1}^{K} N\left(\tau^{(k)}; \tau_0, \eta_0^{-1}\right)$$

where $\tau_0 = 1$ ($\tau_0 = 100$ for the experimental data) and $\delta_0 = 10^{-5}$. In the same way, independent weakly informative Gamma prior distributions with identical shape and scale parameters ($u_0$ and $v_0$) were assume on the precision of the duration of each state

$$p(\rho) = \prod_{k=1}^{K} p\left(\rho^{(k)} \middle| u_0, v_0\right) = \prod_{k=1}^{K} Gam\left(\rho^{(k)}; u_0, v_0\right)$$

where $u_0 = 0.001$ and $v_0 = 1000$, in order to achieve non-informative priors.

###### *2.2.1.2 Posterior updates for the parameters*

Conjugacy and independence of the priors induce variational posterior densities of the same functional form, which are also factorised over states, so that

$$q(\tau) = \prod_{k=1}^{K} q\left(\tau^{(k)}\right) = \prod_{k=1}^{K} N\left(\tau^{(k)}; \hat{\tau}^{(k)}, \hat{\eta}^{(k)^{-1}}\right)$$

Where the posterior mean and precision of the duration of state $k$ are given by

$$\hat{\tau}^{(k)} = \left( \eta_0 \tau_0 + \hat{u}^{(k)} \hat{v}^{(k)} \sum_{d=1}^{D} \gamma_d(k) d \right) \left( \eta_0 + \hat{u}^{(k)} \hat{v}^{(k)} C^{(k)} \right)^{-1}, \qquad \forall k$$
$$\hat{\eta}^{(\mathrm{k})} = \eta_0 + \hat{u}^{(k)} \hat{v}^{(k)} C^{(k)}$$

Similarly, the variational densities of the precision of the durations have the same functional form as their priors and also factorise over states, so that

$$q(\rho) = \prod_{k=1}^{K} q\left(\rho^{(k)}\right) = \prod_{k=1}^{K} Gam\left(\rho^{(k)}; \hat{u}^{(k)}, \hat{v}^{(k)}\right)$$

Where the posterior shape and scale parameters of the posterior Gamma distribution associated with state $k$ are given by

$$\hat{u}^{(k)} = u_0 + \frac{1}{2} C^{(k)}$$

$$\hat{v}^{(k)} = \left( v_0^{-1} + \frac{1}{2} \sum_{d=1}^{D} \gamma_d(k) \left( \left(d - \hat{\tau}^{(k)}\right)^2 + \hat{\eta}^{(k)^{-1}} \right) \right)^{-1}, \qquad \forall k$$

In the above expressions, $\gamma_d^{(k)}$ and $C^{(k)}$ are derived from the sufficient statistics obtained in the VB-E step of the VB algorithm

$$\gamma_d(k) = \sum_{t=1}^{T} \sum_{\substack{i=1 \\ i \neq k}}^{K} \xi_t(i, 1, k, d)$$

$$C^{(k)} = \sum_{d=1}^{D} \gamma_d(k)$$

##### 2.2.2 Log-normal duration model

Similarly to the Normal case, we impose independent conjugate and weakly informative prior distributions on the parameters of the Log-normal distribution associated with each state. Given that the conjugate priors for the parameters of the Log-normal distribution are the same as for the Normal distribution, the expressions for the priors and their variational posteriors are the same as for the Normal case, but replacing the mean and precision of the Normal model with the first and second parameters, respectively of the Log-normal model. In this case, the values of the parameters of the priors (hyperparameters) were $\tau_0 = 4.5$, $\delta_0 = 0.2$, $u_0 = 0.001$ and $v_0 = 1000$.

#### 2.3 Transition model

From the definition of HSMM, the transitions between state segments are governed by a Markov process without self-transitions. Therefore, the transition distribution between states is Multinomial. We then use the usual conjugate priors and assumptions as in the standard HMM (Beal, 2003).

##### 2.3.1 Prior distribution for the transition probabilities

Using standard results from the Bayesian inference of HMM, we assume that the rows $a_{i:}$ of the transition matrix are independent and identically distributed a priori. We then choose the same conjugate non-informative prior distribution for all $a_{i:}$, which is a symmetric non-informative Dirichlet distribution concentration parameter $\alpha_0$ and zero mass on self- transitions,

$$p(A) = \prod_{i=1}^{K} p(a_{i:})$$

$$= \prod_{i=1}^{K} Dir(a_{i1}, \dots, a_{ii-1}, a_{ii+1} \dots, a_{iK}; \alpha_0, \dots, \alpha_0, \alpha_0, \dots, \alpha_0)$$

where $\alpha_0 > 0$. A non-informative prior is achieved by using $\alpha_0 = 1$.

##### 2.3.2 Posterior updates for the transition probabilities

Due to the conjugacy of the prior, the variational posterior of all the rows are Dirichlet, so that

$$q(A) = \prod_{i=1}^{K} q(a_{i:})$$
$$= \prod_{i=1}^{K} Dir(a_{i1}, \dots, a_{ii-1}, a_{ii+1} \dots, a_{iK}; \hat{\alpha}_{i1}, \dots, \hat{\alpha}_{ii-1}, \hat{\alpha}_{ii+1}, \dots, \hat{\alpha}_{iK})$$

with posterior concentration parameters

$$\hat{\alpha}_{ij} = \alpha_0 + \sum_{t=2}^{T} \sum_{d=1}^{D} \xi_t(i, 1, j, d), \qquad \forall i \neq j$$

#### 2.4 Initial probabilities

Similar to the transition distribution, the initial probability distribution $\pi$ is also Multinomial and therefore can be treated in a similar way as one row of the transition matrix.

##### 2.4.1 Prior distribution for the initial probabilities

In the general case (no simplifying initial condition) a conjugate symmetrical Dirichlet prior can be imposed on the initial joint state $(s_1, r_1)$ as

$$p(\pi) = Dir(\pi_{11}, \dots, \pi_{1D}, \pi_{21}, \dots, \pi_{KD}; \gamma_0, \dots, \gamma_0, \gamma_0, \dots, \gamma_0)$$

where $\gamma_0 > 0$. A non-informative prior is implemented using $\gamma_0 = 1$. Under the simplifying initial condition $r_1 = 1$ (so that $\pi_{kd} = 0$ for $d \neq 1$) then the prior can be reduced to

$$p(\pi) = Dir(\pi_{11}, \dots, \pi_{K1}; \gamma_0, \dots, \gamma_0, \gamma_0, \dots, \gamma_0)$$

##### 2.4.2 Posterior updates for the initial probabilities

Due to the conjugacy of the prior, the variational posterior of initial probabilities is also Dirichlet

$$q(\pi) = Dir(\pi_{11}, \dots, \pi_{1D}, \pi_{21}, \dots, \pi_{KD}; \hat{\gamma}_{11}, \dots, \hat{\gamma}_{1D}, \hat{\gamma}_{21}, \dots, \hat{\gamma}_{KD})$$

With concentration parameters

$$\hat{\gamma}_{kd} = \gamma_0 + \gamma_1(k, d), \qquad \forall k, d$$

And under the simplified initial condition

$$\hat{\gamma}_{k} = \gamma_0 + \gamma_1(k, 1), \qquad \forall k$$
